# Improving Causal Gene Identification Using Large Language Models

**DOI:** 10.64898/2026.03.08.710344

**Authors:** Dan Ofer, Hadasa Kaufman

## Abstract

Genome-Wide Association Studies (GWAS) have successfully identified numerous loci associated with complex traits and diseases, yet pinpointing causal genes remains a significant challenge. The reliance on simple proximity-based heuristics is often insufficient due to linkage disequilibrium, gene interactions, and regulatory effects. Recent advancements in Large Language Models (LLMs) have demonstrated potential in automating causal gene identification, but their effectiveness remains limited by knowledge representation and retrieval mechanisms. This study builds on previous research by evaluating LLMs for causal gene identification, with a focus on enhancing performance through Retrieval-Augmented Generation (RAG) and the incorporation of genomic distance information. We replicate prior results using smaller model Qwen2.5—assessing their predictive accuracy using a benchmark dataset from Open Targets. We improved the preformences when integrating RAG-based literature retrieval (F1 = 0.795) and gene distance information (F1 = 0.806). However, the combined approach yielded diminishing returns, suggesting interactions between these enhancements. Error analysis revealed that genomic distance features improved predictions by reinforcing established heuristics, while RAG enhanced domain knowledge but occasionally led to semantic biases. These findings highlight the potential of hybrid approaches in leveraging both structured genomic features and unstructured textual data.

## I. Introduction

Genome-Wide Association Studies (GWAS)[1] have been instrumental in identifying loci associated with various complex traits and diseases, providing a crucial reference point for understanding the contributions of genetics to health[2]. However, since the majority of these loci are located in non-coding regions and contain numerous candidate variants in linkage disequilibrium, it became difficult to precisely pinpoint the specific causal variants and genes[3]. Adding to the challenge the complexity of population genetics, the three-dimensional structure of chromatin, genetic interactions, and genetic linkage[2]. These factors complicate the distinction between mere associations and true causal relationships. Experimental methods, such as ChIP-seq[4], RNA-seq[5] and CRISPR-Cas9[6] often struggle to handle the vast amounts of data and the intricate web of genetic interactions. This bottleneck in driver genes identification underscores the need for additional approaches to effectively translate GWAS findings into actionable biomedical insights.

The naive approach to finding the causal gene is to select the gene closest to the position of the associated variant[7]. Previous attempts addressed this problem by integrating GWAS data with gene expression data to find the affected gene[8]. In recent years, machine learning models have been used to predict causality based on genetics and functional genomics features and outperform the naive distance-based model[9], [2].

Recent advances in the use of Large Language Models (LLMs), such as GPT-3.5 and GPT-4, have shown promise in automating scientific analysis. Recent research has highlighted their effectiveness in performing various biomedical tasks[10], such as summarizing gene functions [11], answering medical queries [12], annotating cell types [13], and identifying causal genetic factors from experimental data in mice [14].

Another challenge in causal gene prediction using LLMs can be the paralog genes. Paralog is a gene that arises from a duplication event within the same genome, leading to two or more genes with similar sequences and potentially overlapping functions[15]. While paralogs can evolve distinct roles over time, their sequence similarity often causes confusion in computational gene prediction. Large Language Models (LLMs) trained on genomic or biomedical texts may struggle with paralogs because they rely on textual patterns and known associations, which can lead to biases.

The study by *Large Language Models Identify Causal Genes in Complex Trait GWAS* [16] demonstrated the potential of LLMs in identification of causal genes, with SOTA results. However, several avenues for improvement remain unexplored.

Specifically advanced techniques such as Retrieval-Augmented Generation (RAG),the integration of domain-specific knowledge (e.g., genomic distance and gene interactions), or more precise prompting strategies could enhance the LLMs performance in this task[17].

This proposal aims to replicate the original results from the aforementioned study[16], to introduce several enhancements and to evaluate them, for the task of improved identification of causal genes. These improvements will be evaluated based on accuracy, and efficiency, with a focus on practical applicability to large-scale datasets.

## II. Problem Formulation

Given a list of candidate genes in the region of associated loci with disease, we want to identify which gene has a causal relationship with the disease.

## III. Datasets

The following datasets will be used in this study:

- **Open Targets gold standard dataset**: The OpenTargets dataset is based on GWAS loci for which there is high confidence (through functional criteria) in the causal gene from *OpenTargets*. We will download the same dataset used in the study: https://github.com/opentargets/genetics-gold-standards/
- **MedRag - Biomedical literature**: A large text dataset of biomedical literature (including 30M PubMed abstracts, Medical textbooks, Wikipedia and Statpearls), used in the MedRAG [18] framework (*Bench-marking Retrieval-Augmented Generation for Medicine*) to augment the LLM’s knowledge base and to reduce hallucinations.
- **Genomic Features**: Data on genomic distances, including the proximity of candidate genes to GWAS derived mutations.

## IV. Proposed Approach

### A. Replication of Baseline Model

We will first replicate the results from *Large Language Models Identify Causal Genes in Complex Trait GWAS* using the same GWAS datasets and methods outlined in the original study. This will serve as a baseline for comparison with the enhanced methods.

### B. Evaluation

Evaluation will be identical to the original paper, with accuracy over the open targets gold dataset (and optionally the temporal validation dataset, if available), using 0-shot approaches, compared to human gold standard annotations as the ground truth.

### C. Enhancements to the Model

Several enhancements will be implemented to improve the performance of the LLM in causal gene identification:

- **Retrieval-Augmented Generation (RAG)**: We will integrate the MedRAG framework [18] to augment the LLM’s knowledge with relevant biomedical literature. This will allow the model to incorporate the latest research and experimental findings that might not be captured in its internal knowledgebase.
- **Genomic Distance as a Feature**: Domain-specific features, such as the genomic distance between candidate genes and the variants, will be included to provide the model with additional context.
- **Improved Prompting Strategies**: We will experiment with more structured and directive prompting, such as chain-of-thought prompting to guide the LLM through a more logical reasoning process that is more aligned with causal inference.

## V. Methods

### A. Pre-processing

The gold standard data downloaded from Open Target with 851 rows of SNPs found associated with a phenotype, with positions, high confidence matched gene as label and list of candidate genes, available from the original authors on Zenodo: https://zenodo.org/records/11391053, [19]. The distances of candidate genes from the associated SNP were calculated on the basis of the ensembl data. For the distance input approaches, the sorted candidate genes and their relative distances from the associated SNP are added the the prompt, in ascending order. After removing duplicated rows and pre-processing and removing duplicated rows,580 rows were left for analysis and evaluation. Each row is a unique phenotype, mutated position and a set of candidate genes near the candidate (in the same locus).

### B. Prediction

We evaluated The llama-3.1 (8B), phi-4 (14B) and Qwen2.5 (32B) models [20], [21], in a 0-shot setting. We also tried supervised, non llm approaches (not shown, includiong sentence transformers), aimed at classifying each combination of target and candidate gene, but abandoned this for computational reasons after observing how performant the locus level prediction setting is. Our prompts used Chain of Thought (COT) approach [22] as shown in Figure-1, We experimented with three Large Language Models (all pretrained Transformers, using 0-shot inference) of varying sizes:

**Figure 1.**
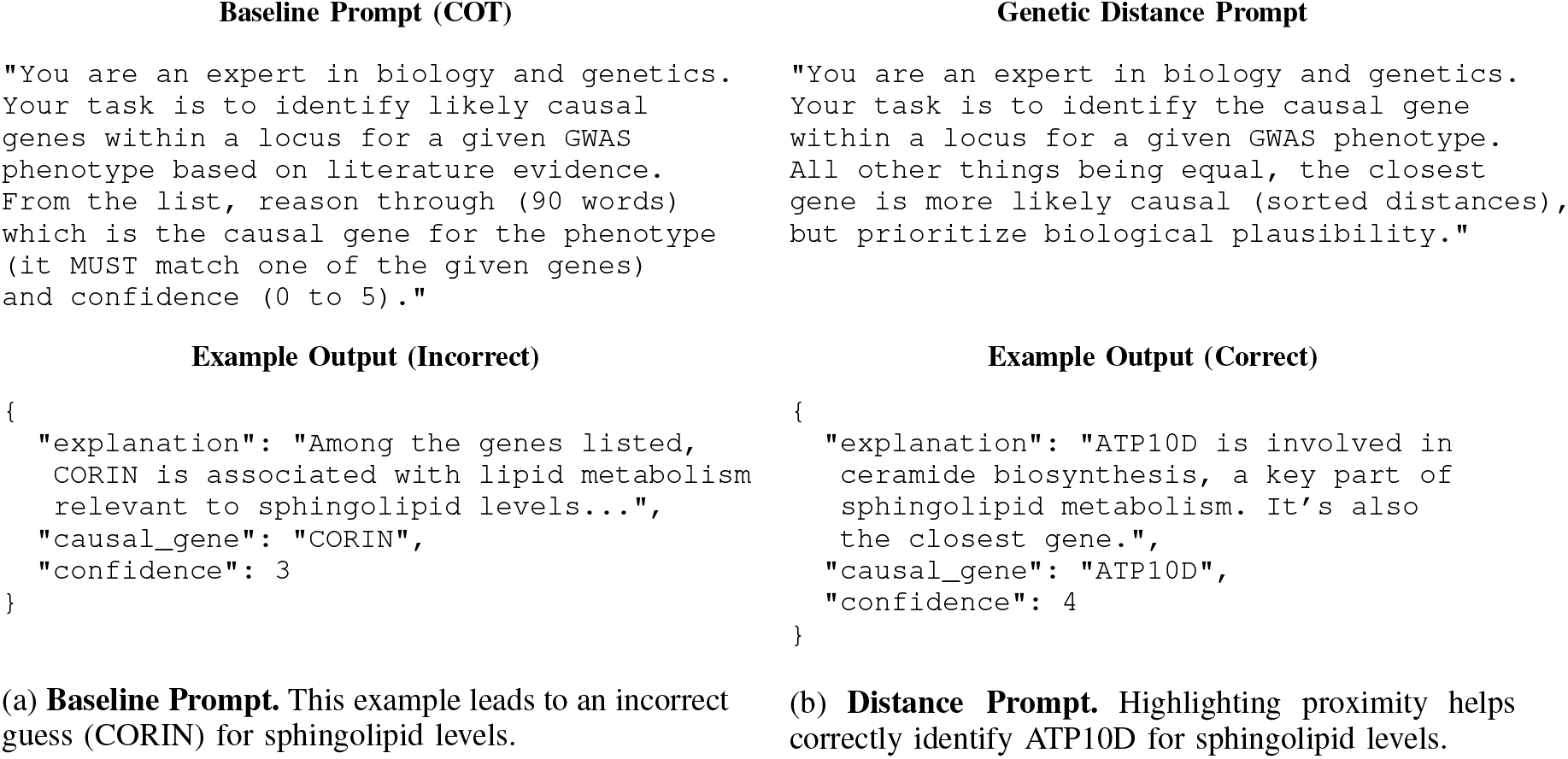
Comparison of Baseline vs. Distance-Based Prompts. Subfigure (a) uses a general chain-of-thought prompt and incorrectly selects `CORIN`. Subfigure (b) integrates sorted gene-distance information and highlights `ATP10D` as the true causal gene for sphingolipid levels. Final suffix of prompts enforcing json structure removed for clarity. Inputs detailed in V-A

- **Llama-3.1 (8B)**: An updated version of the Llama family, containing approximately 8B parameters. It is trained on a diverse corpus of text data. We switched to larger models for our final experiments, after observing that the larger the model used, the better the results. Results not reported.
- **phi-4 (14B)**: A 14B-parameter model described in [20]. phi-4 is trained with data focused on high quality and advanced reasoning.
- **Qwen2.5 (32B)**: A larger, 32B-parameter model introduced by [21]. Although details on its full training dataset remain proprietary, it is reported to have strong capabilities in domain-specific question answering. It was also the largest model we could run.

We focus on the Qwen 32B model for experiments with different inputs (Fig-2), due to its having the highest baseline performance and high prompt adherence (not shown).

**Figure 2.**
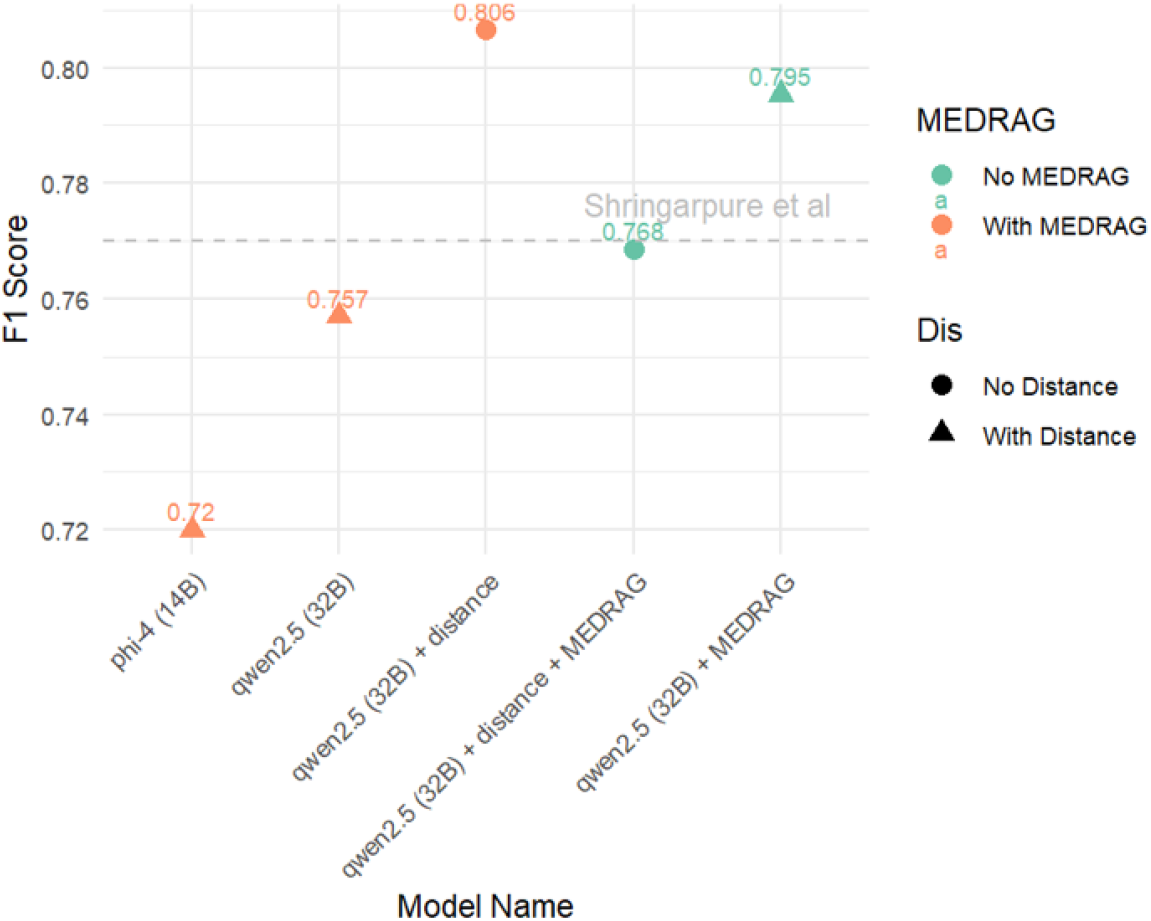
Comparison of model performance. We compare the results of the small phi-4 model with those of the qwen2.5 model. Light blue indicates the use of medRAG and pink indicates a model without RAG. A triangle indicates the use of distances for analysis and a circle indicates a model without distance information. The gray dashed line indicates the performance of Shringarpure et al using GPT4o.

### C. Modeling Approaches

#### 1) Baseline / Chain-of-Thought (COT) Model

Our baseline model used a chain-of-thought prompting strategy with a large language model (LLM) (e.g., Llama-3.1, phi-4, or Qwen2.5) to identify causal genes within a locus for a given GWAS phenotype. In this approach, the model is given a general, structured prompt that asks it to reason through the candidate genes and output a prediction along with an explanation and a confidence score. 1. The model relies on its internal knowledge to make predictions. Although it leverages chain-of-thought reasoning to simulate logical deduction, it sometimes may overemphasize literature popularity or semantic associations, leading to mistakes (for example, selecting a gene based on associated pathways rather than the gene’s proximity to the variant).

#### 2) Retrieval-Augmented Generation (RAG)

Here, the baseline LLM is augmented using the MedRAG framework. This method incorporates a retrieval component that searches a large biomedical text corpus (the aforemention MedRAG corpus) to fetch relevant literature and concatenates it to the prompt/LLMs’ input. We used 25 top documents, retrieved using BM-25. WE note that using a more sophisticated retrieval and reranking system and more items may improve results, but we lacked the computational resources (especially given the large size of the corpus at over 23M+ texts).

#### 3) Gene Distance as a Feature

This approach explicitly including genomic distance information between candidate genes and the associated GWAS SNP. The input genes are also provided, sorted by their distance, with their relative distance from the mutation calculated (rather than using absolute genomic position). This allows leverageing of a well-established heuristic from genetic epidemiology that the closest gene to an associated variant is often—but not always—the causal one. The prompt or the input context is modified to mention that “all other things being equal, the closest gene is more likely causal,” thereby biasing the model toward considering physical proximity as an important factor, as well as biological plausibility or other considerations.

##### a) Combined Approach: RAG + Gene Distance

Finally, both enhancements were combined, with RAG and gene distance given as inputs, using the default prompt.

### D. Model evaluation

For each of the models F1 score was calculated using the following formula:

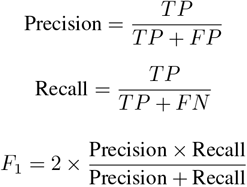

The F1 score of all models was compared with the performance of Shringarpure et al, including their approach. Specifically, “the predictions from each method were converted to 0/1/NA, with 1 assigned if the method made a prediction and it matched the annotated causal gene”, and 0 assigned if the method made a prediction that did not match the annotated causal gene (including malformed prompts or genes not in the candidate list - of which we had 0 when using the 14B+ models).

## VI. Results

The performance of the models showed improved F-score with increasing model size. For a 32B model, the addition of medRAG improved the performance to an F-score of 0.795, and the addition of information on gene distances and sorting by distance from the SNP associated with the phenotype improved it even further to an F-score of 0.806. Interestingly, the combination of the two upgrades together produced a model that was less good than the model with each upgrade alone, although the model was still better than a model with no additions at all (Figure 2).

### A. Error analysis

For the basic model and the RAG and distance-based model, we distinguished between errors where the model predicted a paralog for the correct gene and cases where the gene the model predicted as causal was a completely different gene - Fig-3. For several cases where different models gave different results, we examined the explanations that the models gave to characterize the capabilities and limitations of each model. We provide some examples in the appendix - A.

**Figure 3.**
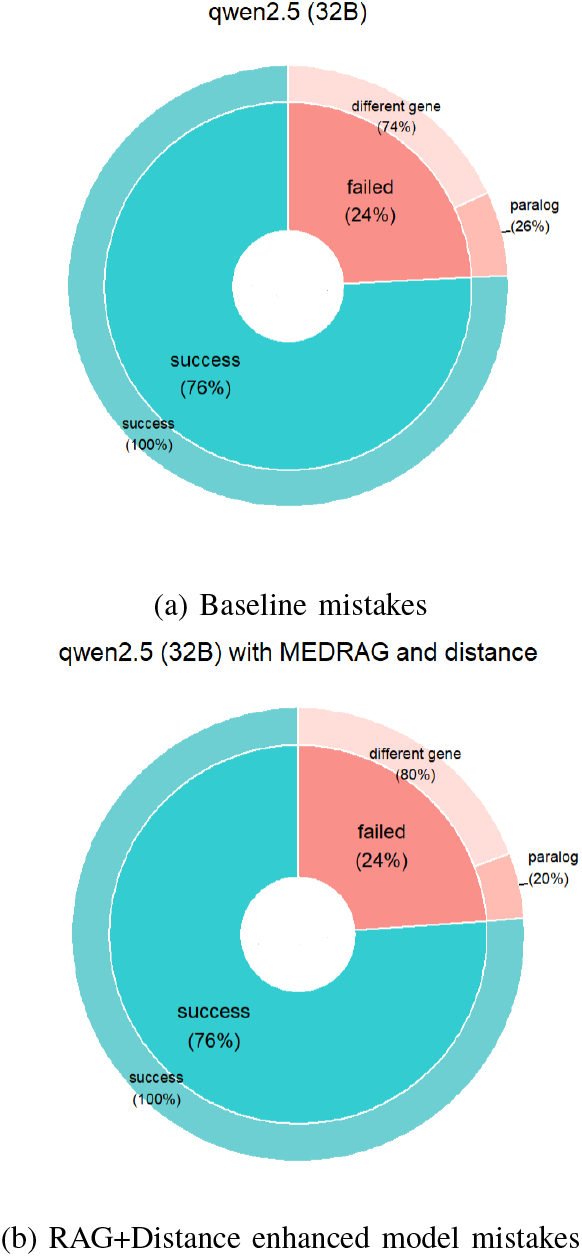
Mistake type distribution between (a) baseline model, and (b) with additional literature and genetic distance information. We note the difference in mistaken paralogs

The base model with COT and the model with MedRAD and distances predict the same gene in 76% of the cases. 90.1% of them were true predictions and 9.9% failed with the same gene. The entire 26% predictions was different between the models, while 39.8% of them successed in the model with MedRAD and distances only, 32.6% successed in the base model only and 27.6% feiled in both with dieewrwnt false prediction.

For example, when the LLM model has access to genomic distance data, it correctly predicts ATP10D as the causal gene, explaining that ATP10D is involved in ceramide biosynthesis and trafficking, directly linking it to sphingolipid metabolism. The genetic study referenced further supports its role in influencing sphingolipid levels, making it a strong candidate for causality.A

In contrast, when the model does not utilize distance information, it incorrectly predicts CORIN as the causal gene. While CORIN is associated with lipid metabolism, there is no direct evidence linking it specifically to sphingolipid levels, demonstrating a weaker causal link.

## VII. Discussion

In our work, we show performance that matches or exceeds that of the original paper, without any meaningfully increased complexity. We see improved performance as being easily reachable, using larger models (e.g. 70B, or gpt 4o or O1/O3-mini), especially given the overall performance trend not saturating. An important caveat is that we did not have access to the temporal test set of Weeks et al, mentioned in [16], and this matters due to the unknown risk that information about the target labels may have been readable by the models in their pretraining phase. The cutoff of the Qwen-32 model is also not known. A hidden test set could help with this, along with any novel candidates for research. Still, we believe that, given past standards [16], this is still a promising and likely generalizable result, given the importance of this task, the generalization shown in the original paper and the fact that the applications of this model would be as a decision aid.

In our analysis of the types of mistakes made (e.g. paralogs fig-3) and manual analysis of model explanations (with many relying on “strong literature evidence”) explanations indicates that literature evidence acted as to confuse the models. When the model relied more on data and known biological facts, including distance, i.e., a more “data driven” approach, its performance is better and relies more on what the gene does (Causality) than popular genes or variants. We note also that the annotators of the gold-standard data may rely on genetic distance as part of their decision making process, and making it available may also explain the models improved “alignment” with human decisions.

## Appendix A. Distance+Rag improved predictions

We show here some examples of genes whose predictions were mistaken with the original model, but had a correct prediction by the final model (using RAG and gene distance). In total, there were 48 such cases.

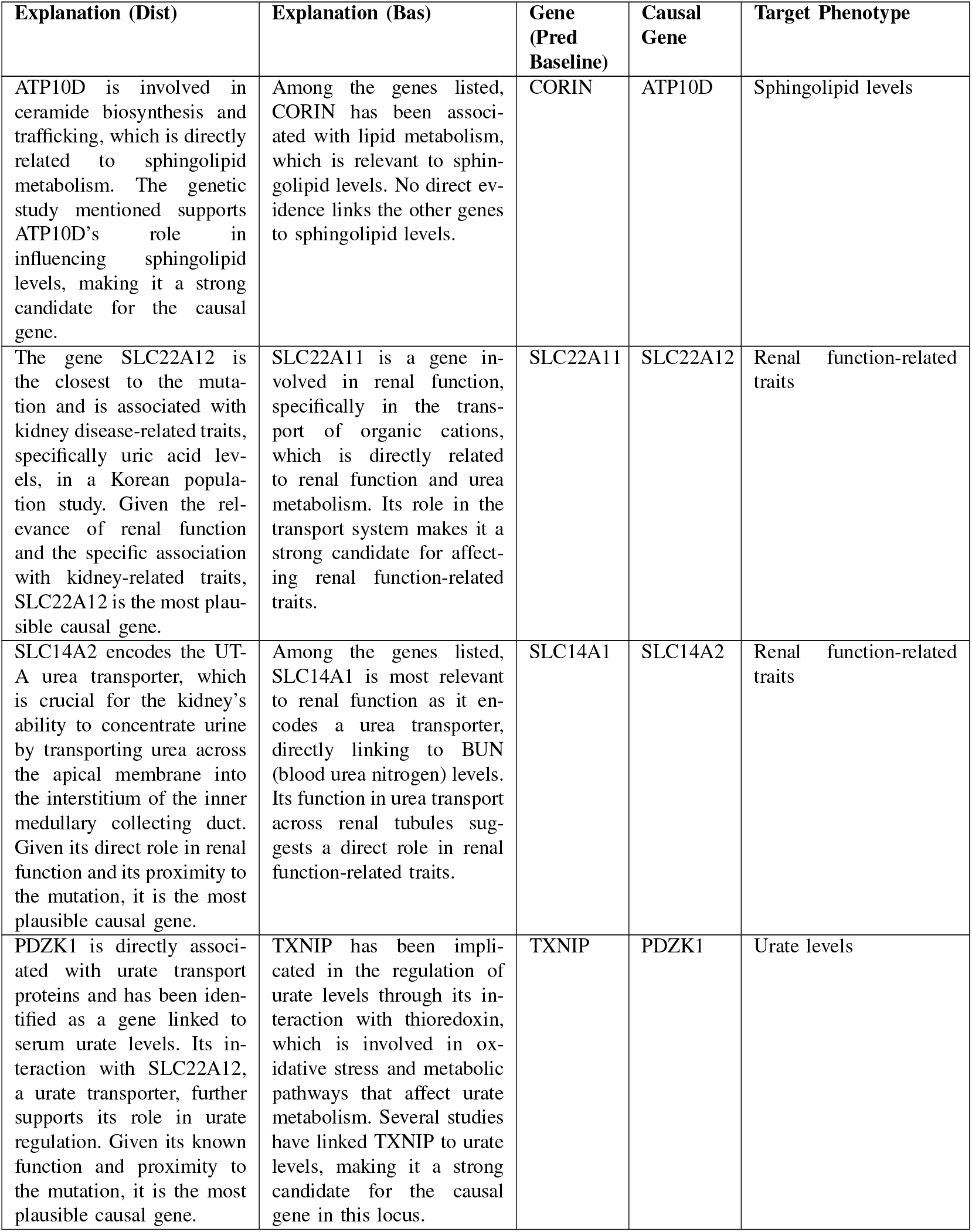
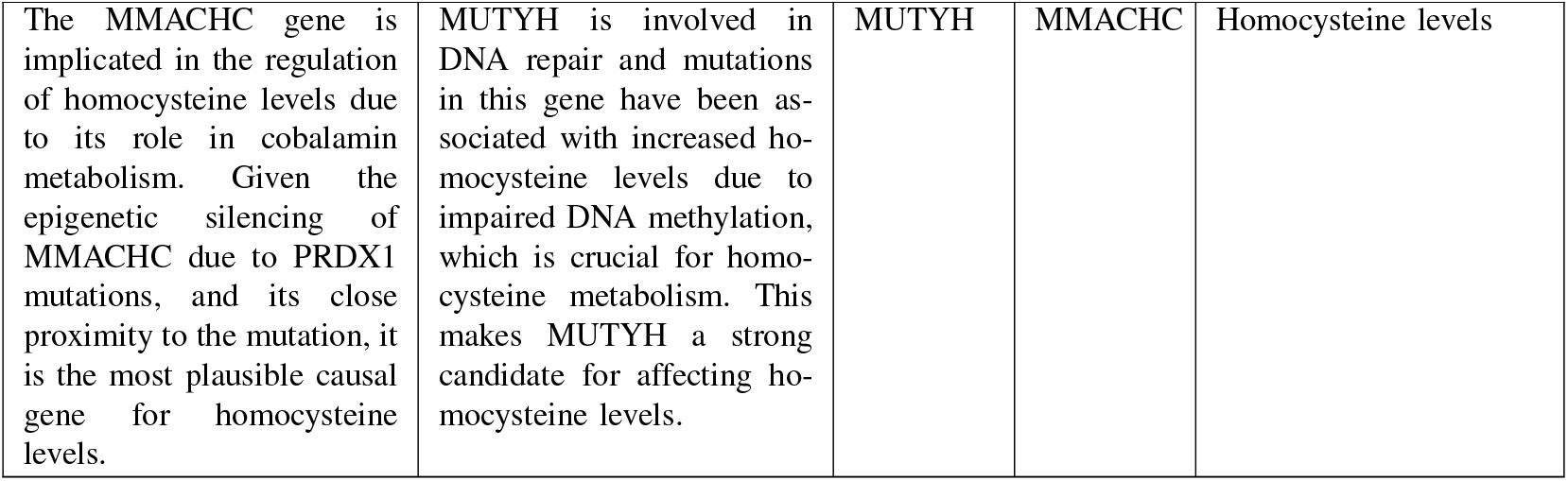

| Explanation (Dist) | Explanation (Bas) | Gene (Pred Baseline) | Causal Gene | Target Phenotype |
| --- | --- | --- | --- | --- |
| ATP10D is involved in ceramide biosynthesis and trafficking, which is directly related to sphingolipid metabolism. The genetic study mentioned supports ATP10D's role in influencing sphingolipid levels, making it a strong candidate for the causal gene. | Among the genes listed, CORIN has been associated with lipid metabolism, which is relevant to sphingolipid levels. No direct evidence links the other genes to sphingolipid levels. | CORIN | ATP10D | Sphingolipid levels |
| The gene SLC22A12 is the closest to the mutation and is associated with kidney disease-related traits, specifically uric acid levels, in a Korean population study. Given the relevance of renal function and the specific association with kidney-related traits, SLC22A12 is the most plausible causal gene. | SLC22A11 is a gene involved in renal function, specifically in the transport of organic cations, which is directly related to renal function and urea metabolism. Its role in the transport system makes it a strong candidate for affecting renal function-related traits. | SLC22A11 | SLC22A12 | Renal function-related traits |
| SLC14A2 encodes the UT-A urea transporter, which is crucial for the kidney's ability to concentrate urine by transporting urea across the apical membrane into the interstitium of the inner medullary collecting duct. Given its direct role in renal function and its proximity to the mutation, it is the most plausible causal gene. | Among the genes listed, SLC14A1 is most relevant to renal function as it encodes a urea transporter, directly linking to BUN (blood urea nitrogen) levels. Its function in urea transport across renal tubules suggests a direct role in renal function-related traits. | SLC14A1 | SLC14A2 | Renal function-related traits |
| PDZK1 is directly associated with urate transport proteins and has been identified as a gene linked to serum urate levels. Its interaction with SLC22A12, a urate transporter, further supports its role in urate regulation. Given its known function and proximity to the mutation, it is the most plausible causal gene. | TXNIP has been implicated in the regulation of urate levels through its interaction with thioredoxin, which is involved in oxidative stress and metabolic pathways that affect urate metabolism. Several studies have linked TXNIP to urate levels, making it a strong candidate for the causal gene in this locus. | TXNIP | PDZK1 | Urate levels |
| The MMACHC gene is implicated in the regulation of homocysteine levels due to its role in cobalamin metabolism. Given the epigenetic silencing of MMACHC due to PRDX1 mutations, and its close proximity to the mutation, it is the most plausible causal gene for homocysteine levels. | MUTYH is involved in DNA repair and mutations in this gene have been associated with increased homocysteine levels due to impaired DNA methylation, which is crucial for homocysteine metabolism. This makes MUTYH a strong candidate for affecting homocysteine levels. | MUTYH | MMACHC | Homocysteine levels |

